# The Positive Switching RSFP Padron2 Enables Live-Cell RESOLFT Nanoscopy Without Sequential Irradiation Steps

**DOI:** 10.1101/2020.09.29.318733

**Authors:** Timo Konen, Tim Grotjohann, Isabelle Jansen, Nickels Jensen, Stefan W. Hell, Stefan Jakobs

**Affiliations:** Department of NanoBiophotonics, Max Planck Institute for Biophysical Chemistry, Göttingen, Germany; Department of Neurology, University of Göttingen, Göttingen, Germany

**Keywords:** Super-resolution microscopy, Padron, switching, live cell, fluorescent protein

## Abstract

Reversibly switchable fluorescent proteins (RSFPs) can be repeatedly transferred between a fluorescent on- and a non-fluorescent off-state in response to irradiation with light of different wavelengths. Negative switching RSFPs are switched from the on- to the off-state with the same wavelength which also excites fluorescence. Positive switching RSFPs have a reversed light response where the fluorescence excitation wavelength induces the transition from the off- to the on-state. Reversible saturable optical linear (fluorescence) transitions (RESOLFT) nanoscopy utilizes these switching states to achieve diffraction-unlimited resolution, but so far has primarily relied on negative switching RSFPs by using time sequential switching schemes.

Based on the green fluorescent RSFP Padron, we engineered the positive switching RSFP Padron2. Compared to its predecessor, it can undergo 50-fold more switching cycles while displaying a contrast ratio between the on- and the off-state of more than 100:1. Because of its robust switching behavior, Padron2 supports a RESOLFT imaging scheme that entirely refrains from sequential switching as it only requires beam scanning of two spatially overlaid light distributions. Using Padron2, we demonstrate live-cell RESOLFT nanoscopy without sequential irradiation steps.

## Introduction

Nanoscopy, or diffraction-unlimited super-resolution fluorescence microscopy, enables the visualization of cellular structures at the nanoscale. The key to fundamentally overcome the diffraction barrier is to make adjacent molecules discernible through a fluorescence on/off state transition forcing adjacent fluorophores to emit sequentially.^1^ This separation of adjacent fluorophores can be implemented either in a coordinate-targeted or in a coordinate-stochastic way (for reviews see ^2, 3^).

RESOLFT (reversible saturable optical linear (fluorescence) transitions) nanoscopy is a coordinate-targeted approach that relies on reversibly switchable fluorophores.^4–7^ In point-scanning RESOLFT nanoscopy, a single laser beam creating a doughnut shaped intensity distribution with a zero at its center is used to transfer molecules into a non-fluorescent off-state. Thereby, the on-state molecules and hence emission are limited to a central region smaller than the diffraction limit, which is read out by a regularly focused beam.

Most implementations of RESOLFT nanoscopy rely on reversibly switchable fluorescent proteins (RSFPs), which belong to the group of GFP-like fluorescent proteins.^8, 9^ These proteins feature a beta-barrel structure with a central alpha-helix containing the autocatalytically formed chromophore.^10^ RSFPs can be reversibly toggled between a fluorescent on- and a non-fluorescent off-state by irradiation with two different wavelengths. Because RSFPs are metastable in both the on- and the off-state and the quantum yield for switching is comparatively high, the light doses and intensities required for overcoming the diffraction barrier are low compared to basically all other nanoscopy approaches.^11^ In fact, the light intensities used are similar to those applied in live-cell confocal fluorescence microscopy. As the intensity of irradiation is an important factor that determines phototoxicity^12, 13^, RESOLFT nanoscopy is particularly suitable for live-cell recordings.

The switching behavior of most RSFPs has been classified as either negative or positive switching.^14^ Negative switching RSFPs are switched from the on- to the off-state with the same wavelength which is also used for fluorescence excitation. In positive switching RSFPs the excitation wavelength induces the off-to-on transition. In both classes of RSFPs, the respective other switching direction is triggered by a second, shorter wavelength. At present, almost all RESOLFT implementations rely on RSFPs with a negative switching mode.^9^ In a typical RESOLFT scheme using negative switching RSFPs, fluorophores are switched sequentially:^4, 5, 11, 15–19^ After initial switching to the on-state, RSFPs are switched off with a doughnut-shaped beam or a standing wave light pattern and remaining on-state fluorophores are probed with a regularly focused beam. Because for negative switching RSFPs fluorescence readout and off-switching are triggered by the same wavelength, central fluorophores are switched off during readout. As a consequence, the switching and readout sequence often needs to be repeated in order to collect enough photons if expression levels are low. This procedure is unfavorable as it increases the image acquisition time and the light dose applied to the sample.

Positive switching RSFPs can be used to overcome the problem of limited fluorescence readout per switching cycle because fluorescence excitation triggers the on-switching process and hence the proteins in the center of the doughnut can be kept in the on-state for an arbitrary time during readout. Here, to achieve sub-diffraction resolution, the molecules in the periphery have to be kept in the off-state, which can be achieved by superposition of the regularly focused excitation light with the doughnut-shaped off-switching beam. In principle, the two overlaid beams can be scanned together over the sample to record a super-resolved image, without the requirement for sequential irradiation steps. For simplicity, we refer to this approach as one-step RESOLFT nanoscopy.

In fact, the initial demonstration of RESOLFT nanoscopy 15 years ago was performed in the one-step mode.^7^ However, the utilized protein, asFP595, showed a poor switching performance and is an obligate tetramer. It is therefore not suitable as a fusion tag in live-cell imaging applications. Consequently, following realizations of the RESOLFT concept refrained from positive switching RSFPs und used negative switching RSFPs with sequential irradiation steps due to the unavailability of suitable probes.

Aside from asFP595,^20, 21^ only few positive switching proteins have been reported so far, and none of them display the switching performance required. rsCherry,^22^ a red-emitting RSFP, and Padron,^14^ a green-emitting RSFP engineered from Dronpa,^23^ both display a low resistance to switching fatigue and slow switching kinetics. These characteristics were improved to some extent in Kohinoor,^24^ a variant based on Padron. It has been used for demonstration of point-scanning RESOLFT, but we found that Kohinoor is still prone to switching fatigue. However, resistance against switching fatigue is a key requirement for one-step RESOLFT imaging.

One-step RESOLFT nanoscopy requires a positive switching RSFP which does not emit fluorescence upon irradiation with the off-switching wavelength. In addition, the vast majority of peripheral RSFPs need to reside in the off-state despite being irradiated with on-switching light, which requires the ensemble equilibrium state under simultaneous irradiation to be adjustable freely and quickly by changing the relative light intensities. If the equilibrium is reached fast relative to the beam movement during sample scanning, simultaneous irradiation with both superimposed beams alone should suffice to overcome the diffraction barrier.

To engineer a positive switching RSFP with improved characteristics, we chose to rely on the well-described positive switching protein Padron.^14^ This RSFP displays poor expression at 37 °C and it features slow switching kinetics as well as high switching fatigue. On the other side, Padron displays high molecular brightness and switching contrast. Importantly, several X-ray structures of Padron are available.^25, 26^ In addition, the switching mechanism of Padron has been investigated by kinetic crystallography^25, 26^ and ultra-fast spectroscopy,^27, 28^ at cryo-temperatures^29^ as well as with extensive molecular dynamics simulations,^25, 30^ providing a solid data base for a semi-rational engineering approach.

We generated the new positive switching RSFP Padron2, which outperforms all previous positive switching RSFPs for RESOLFT nanoscopy. With Padron2, we established RESOLFT nanoscopy on living cells without the need for sequential switching steps.

## Results

### Development of Padron2

All currently available positive switching RSFPs exhibit severe limitations for their application in RESOLFT nanoscopy. In order to explore possibilities of improvement, we decided to screen for novel positive switching RSFPs based on Padron as a template in a semi-rational approach. Padron is a green fluorescent protein (emission wavelength: 522 nm) and fluorescence can be exited with light of 488 nm wavelength.^14^ This wavelength also transfers the protein from the off-into the on-state. Light of 405 nm wavelength switches the protein from the on-into the off-state. We chose Padron as a template because it has been well characterized, has a high molecular brightness, and can be switched to low ensemble off-state fluorescence intensity.

We performed 16 rounds of error-prone and site-directed saturation mutagenesis on the Padron coding sequence. For site-directed mutagenesis, amino acid positions selected based on crystal structures of Padron switching states (PDB 3LS3, 3LSA, 3ZUJ, and 3ZUL) were addressed. For random mutations, PCR-based error prone mutagenesis was performed. After each round of mutagenesis, plasmid libraries encoding the Padron mutants were transformed into Escherichia coli bacteria and plated on agar plates. In every screening round up to four agar plates with ~1,000 bacterial colonies each were analyzed with an automated fluorescence microscope equipped with laser diodes for the switching of the RSFPs. This enabled repeated switching of the RSFP fluorescence with μs precision and at light intensities typically used in RESOLFT imaging. The fluorescence modulation in response to alternating irradiation with light of 405 nm and 488 nm wavelength allowed for simultaneous determination of switching kinetics, residual off-state fluorescence intensity, switching fatigue and effective brightness of each bacterial colony. After each screening round, the best performing variants were sequenced and chosen for further characterization and mutagenesis.

Finally, we identified a Padron variant with substantially improved properties that differed at 11 positions from its template Padron: M40V, T58S, R66K, A69I, S82L, Y114F, L141P, F173S, S190A, E218G, and R221G (Suppl. Figs. 1, 2). In addition, we modified the N- and C-terminal ends of the protein to match those of EGFP, which had previously been reported to improve the tagging performance of fluorescent proteins.^31^ We named this new RSFP Padron2. To evaluate the properties of Padron2, we systematically compared it to Padron and Kohinoor, which both have previously been used for cellular imaging.

### General properties of Padron2

Padron2 has a fluorescence excitation maximum at 492 nm and an emission maximum at 516 nm (Fig. 1a, Tab. 1). Thus, it is spectrally slightly blue-shifted in comparison to Padron and it is similar to Kohinoor in this respect. The fluorescence lifetime of Padron2 is slightly shorter compared to Padron and Kohinoor (3.0 ns vs 3.4 ns and 3.5 ns) (Tab. 1). At pH 7.5, ~67 % of Padron2 molecules adopted the on-state in equilibrium, while Padron and Kohinoor predominantly resided in the off- or on-state, respectively (Tab. 1; Suppl. Fig. 3a-c). Padron2 molecules in solution displayed two distinct absorption peaks in the on-state at 384 nm and 495 nm (Fig. 1b; Tab. 1), resulting from the protonated and deprotonated form of the chromophore.^10, 32^ In on-state Padron2, compared to Padron, the occupancy of the protonated chromophore was increased, while that of the deprotonated chromophore was decreased (Fig. 1b-d). The lowered absorption of the on-state contributed to the decrease in molecular brightness of on-state Padron2 (7.9) at pH 7.5 compared to Padron (25.6) and was similar to that of Kohinoor (9.2) (Tab. 1). For Kohinoor a molecular brightness of 44.7 was previously reported.^24^ However, this value was determined at an unphysiological pH value of 10.0 and is hence irrelevant for a proper comparison. Padron has a pKa of 5.9 (Tab. 1) and fluorescence decreases strongly at pH values above 7.0 (Fig. 1e). As pH values between 7 and 8 are common in cells, this is an undesirable property. This pH instability is eliminated in Padron2, which is stable at pH values as high as 11.0 (Fig. 1e; Suppl. Fig. 4), with two pKa-values of 6.6 and 9.1 (Tab. 1).

**Figure 1.**
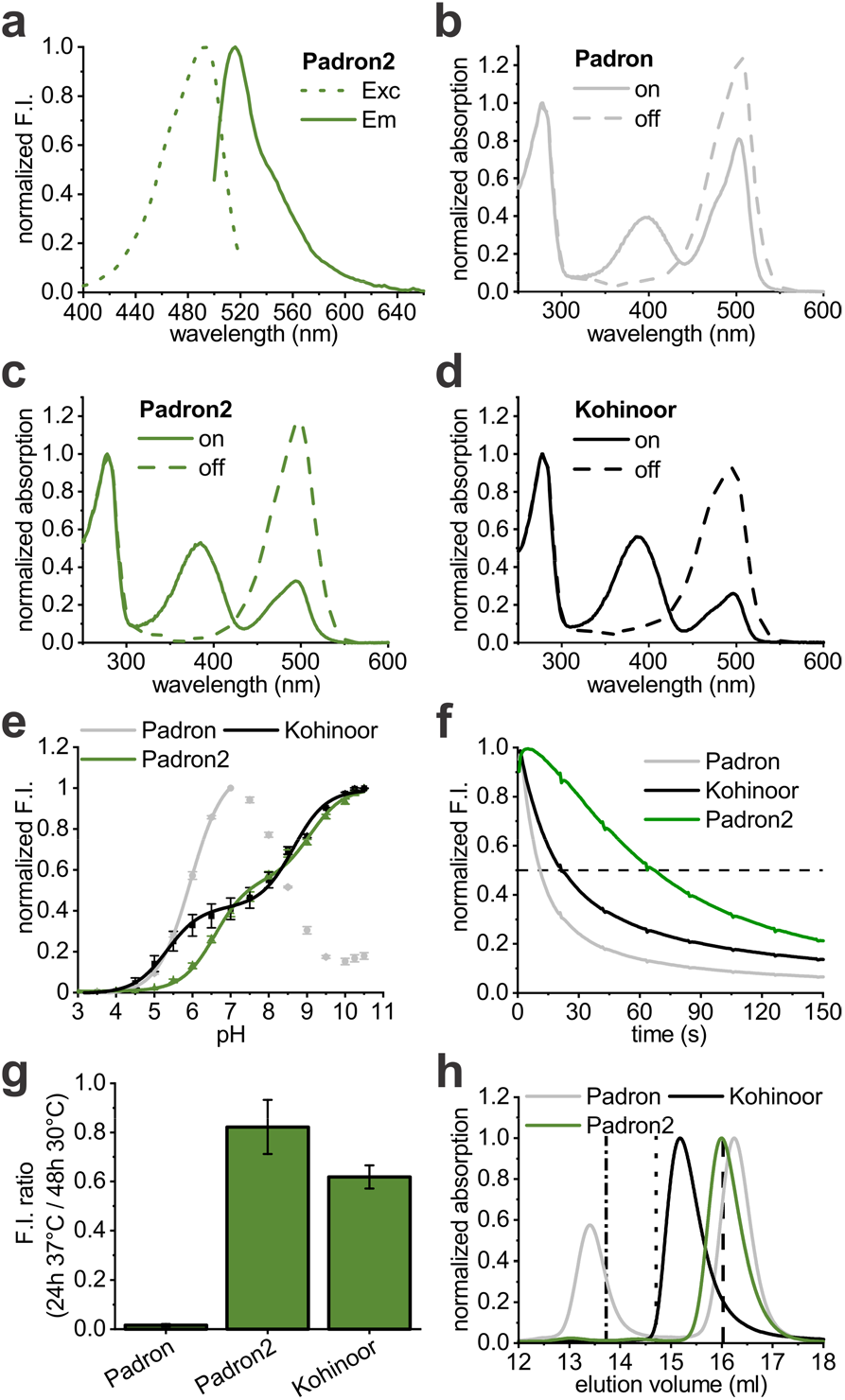
Protein characteristics. **(a)** Excitation (Exc) and emission (Em) spectra of Padron2. **(b, c, d)** Absorption spectra of Padron (b), Padron2 (c), and Kohinoor (d) in the on- and off-state. **(e)** pH-dependent normalized fluorescence intensity of purified protein solutions at the equilibrated state. **(f)** Photobleaching in bacterial colonies under 488 nm irradiation. **(g)** Fluorescence intensity ratios measured in bacterial colonies after 24 h growth at 37 °C and 48 h growth at 30 °C. **(h)** Size exclusion chromatography of purified protein samples. Absorption was measured at 280 nm, vertical lines indicate the peak elution volume of tetrameric DsRed (dashed-dotted line), dimeric dTomato (dotted line), and monomeric mEGFP (dashed line).

**Table 1.** Protein characteristics of Padron, Padron2, and Kohinoor

| Protein | Experimental conditions | Padron | Padron2 | Kohinoor |
| --- | --- | --- | --- | --- |
| **Absorption maximum [nm]** | on-state | 398, 504 | 384, 495 | 388, 496 |
|  | off-state | 505 | 498 | 496 |
| **Excitation maximum [nm]<sup>a</sup>** | equilibrated, pH 7.5 | 502 | 492 | 496 |
| **Emission maximum [nm]<sup>a</sup>** | equilibrated, pH 7.5 | 522 | 516 | 518 |
| **Emission maximum [nm]** | on-state, pH 7.5 | 519 | 513 | 514 |
| **Extinction coefficient [M<sup>-1</sup>cm<sup>-1</sup>]** | pH 7.5, on-state maximum | 39,900 ± 1,500<sup>b</sup> | 16,200 ± 600 | 12,600 ± 400<sup>c</sup> |
| **Quantum yield** | on-state, pH 7.5 | 0.64<sup>d</sup> | 0.49 ± 0.02 | 0.73 ± 0.03<sup>e</sup> |
| **Molecular brightness** | on-state, pH 7.5 | 25.60 | 7.90 | 9.20 |
| **pKa** | equilibrated, in solution | 5.9 | 6.6, 9.1 | 5.3<sup>f</sup>, 8.6 |
| **Time to bleach to 50 % F.I. [s]** | in bacterial colonies | 11.4 | 67.4 | 21.5 |
| **Fluorescence lifetime [ns]** | in solution, pH 7.5 | 3.4 ± 0.02 | 3.0 ± 0.05 | 3.5 ± 0.09 |
| **Equilibrium F.I. [% of on-state]** | in solution, pH 7.5 | 8.9 ± 1.2 | 66.7 ± 2.4 | 96.7 ± 1.8 |
| **Residual off-state F.I. [% of on-state]** |  | 4.0 ± 0.3 | 1.9 ± 0.1 | 2.0 ± 0.1 |
| **Residual off-state F.I. [% of on-state]** | in bacterial colonies | 2.27 ± 0.02 | 1.41 ± 0.09 | 8.42 ± 0.63 |
| **Switching half-time [ms]<sup>g</sup>** | in bacterial colonies | 81.9 | 32.8 | 37.6 |
| **Number of cycles at 50 % maximal intensity** | in bacterial colonies | 19 (113<sup>h</sup>) | 990 (321<sup>i</sup>) | 37 (182<sup>h</sup>) |

<sup>a</sup>for spectra, see Fig. 1a, Suppl. Fig. 3e,f <sup>b</sup>published: 43,000;<sup>14</sup> <sup>c</sup>published: 62,900 ± 136, measured at pH 10; <sup>24</sup> <sup>d</sup>published value;<sup>14</sup> <sup>e</sup>published value: 0.71 ± 0.05;<sup>24</sup> <sup>f</sup>published values: 5.9, 8.6;<sup>24</sup> <sup>g</sup>measured at 1.3 kW/cm<sup>2</sup> 488 nm intensity <sup>h</sup>measured with Padron2 settings; <sup>i</sup>measured with Kohinoor settings. F.I., fluorescence intensity.

An important parameter for the usability of a fluorescent protein in microscopy is its stability against photobleaching. Under continuous irradiation at 2.3 kW/cm^2^ in bacterial colonies, Padron2 displayed a 2- and 6-fold increased resistance to photobleaching in comparison to Kohinoor and Padron (Fig. 1f, Tab. 1), respectively.

Padron exhibits poor expression at 37 °C but is well expressed at lower temperatures. To evaluate the expression of Padron2 at 37 °C, we compared the fluorescence of bacterial colonies grown for 24 h at 37 °C with colonies grown for 48 h at 30 °C (Fig. 1g; Suppl. Fig. 3d). Whereas E. coli colonies expressing Padron2 showed almost the same fluorescence signal under both conditions, Kohinoor exhibited a significantly weaker fluorescence signal upon growth at 37 °C and Padron expressing colonies exhibited almost no mature protein at 37 °C.

Taken together, with regard to key spectroscopic properties used to characterize fluorescent proteins in vitro, Padron2 is similar to Padron and Kohinoor, and with respect to photostability and bacterial expression at 37 °C, it outperforms these RSFPs.

### Tagging with Padron2

In size exclusion chromatography, Padron2 eluted as a monomer (Fig. 1h). In comparison, Padron eluted with two distinct populations as tetramer as well as a monomer, suggesting slow dissociation kinetics.^33^ Kohinoor eluted between the volumes of a dimer and a monomer, indicating fast dissociation and association rates of a dimeric interaction (Fig. 1h) under these conditions.^33^

To analyze the performance of Padron2 as a tag in fusion proteins, we cloned several Padron2 fusion constructs targeted to various cellular structures and expressed them in human HeLa cells (Fig. 2). All Padron2 fusion constructs tested localized correctly. Constructs included fusions to vimentin (VIM), keratin 18 (KRT18), the actin binding peptide lifeact, the microtubule associated protein Map2 (MAP2), the centromere protein C1 (CENPC1), caveolin 1 (CAV1), the nuclear pore complex protein nucleoporin 50 (NUP50) as well as to the histone H2bn (HIST1H2BN). We also targeted Padron2 to the lumen of the ER, the mitochondrial matrix, peroxisomes and the cytosol (Fig. 2).

**Figure 2.**
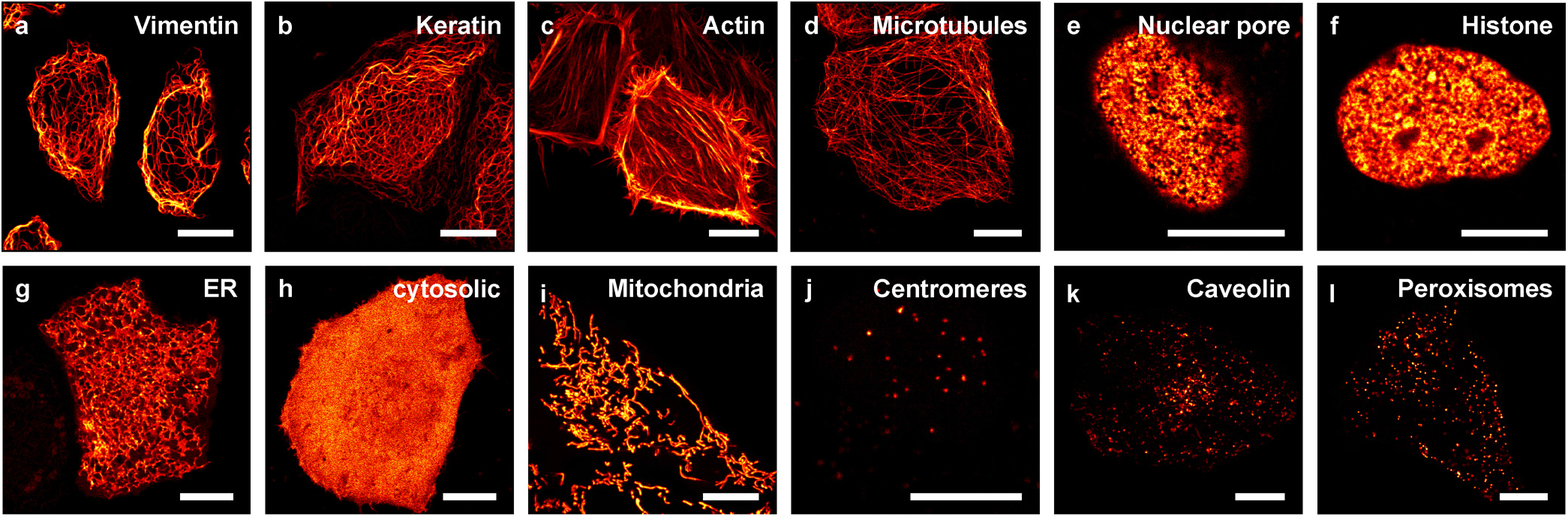
Padron2 fusion constructs in HeLa cells. The fusion proteins were transiently expressed in HeLa cells. **(a)** Vimentin (vimentin-Padron2), **(b)** keratin (keratin18-Padron2), **(c)** actin (lifeact-Padron2), **(d)** microtubules (Map2-Padron2), **(e)** nuclear pore (Padron2-Nup50), **(f)** histone (Padron2-histone H2bn), **(g)** endoplasmic reticulum (TS-Padron2-ER retention signal) **(h)** cytosolic, **(i)** mitochondria (TS-Padron2), **(j)** centromeres (Padron2-CenpC), **(k)** caveolin (caveolin-1-Padron2), **(l)** peroxisomes (Padron2-TS). Images were recorded 24 h post transfection in a single plane (e-h, j-l) or as maximal projection of a z-stack (a-d, i). TS, targeting sequence. Scale bars: 10 μm.

### Switching characteristics of Padron2

In RESOLFT imaging, three switching parameters of an RSFP are of key relevance: 1) The switching kinetics, 2) the residual fluorescence intensity in the ensemble off-state, and 3) the switching fatigue, which is the switching induced photodestruction of the fluorophore. Padron and Kohinoor are particularly poor with respect to switching fatigue, so we had screened for a variant that showed a substantially higher resistance against switching fatigue, while at the same time we had aimed at generating an RSFP that enabled a faster off-switching and a lower residual fluorescence background than Padron.

To compare the switching fatigue of Padron2, Padron, and Kohinoor, we cycled the fluorescence of the respective proteins expressed in E. coli between 5 % residual off-state fluorescence intensity (10 % in case of Kohinoor because it could not be switched below 8 %) and 95 % of the full on-state by choosing appropriate light intensities and irradiation times. The fluorescence of bacterial colonies expressing Padron2 was cycled almost 1000 times before the fluorescence intensity was bleached to 50 %. With Padron and Kohinoor, we achieved only 19 and 37 cycles, respectively (Fig. 3a; Tab. 1). Hence, the final Padron2 displayed an outstanding resistance against switching fatigue.

**Figure 3.**
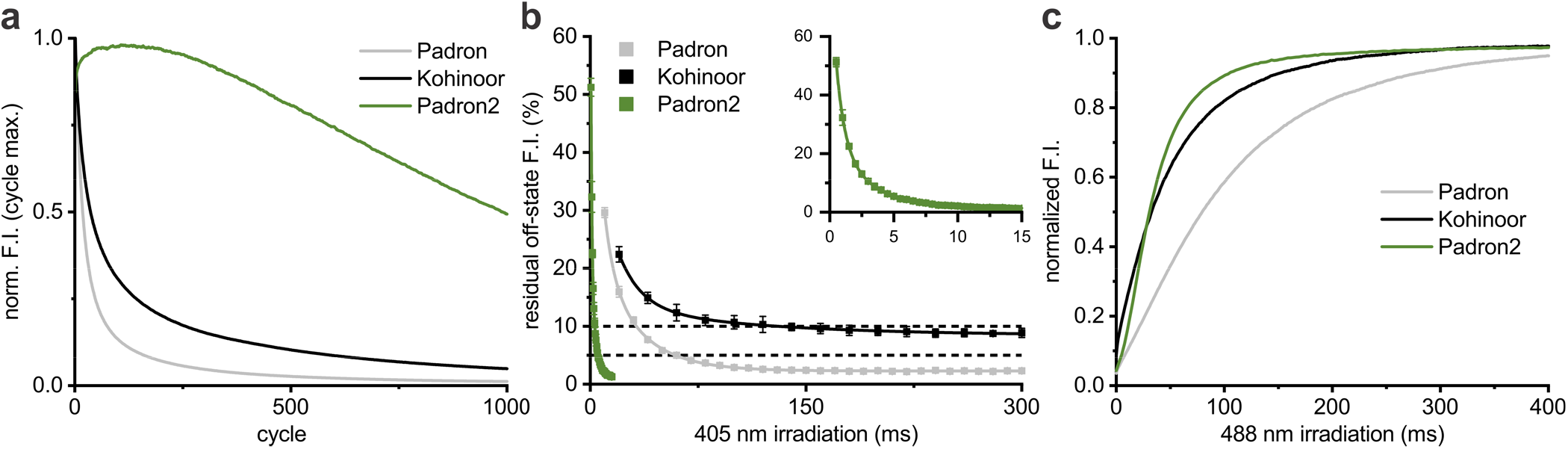
Switching performance **(a)** Switching fatigue in bacterial colonies. Line graphs represent maximal fluorescence intensities of the on-switching curves of every cycle. **(b)** Residual fluorescence intensity in bacterial colonies after different 405 nm irradiation times with an intensity of 4.1 kW/cm^2^. Squares represent averaged measurements, while the line graph is the respective exponential decay fit. Inset graph displays Padron2 data from the main graph with adjusted axis scales. **(c)** Normalized on-switching curves in bacterial colonies. Padron and Padron2 had previously been switched to 5 % residual off-state fluorescence intensity, while Kohinoor had been switched to 10 %.

Padron2 has the distinct advantage of emitting very little fluorescence when being excited by light of 405 nm, the wavelength that induces the transition from the on- to the off-state (Fig. 1a). This advantage, however, renders it also challenging to probe the off-switching speed, as one cannot measure the decrease of fluorescence upon irradiation with 405 nm light. To overcome this challenge, bacterial colonies expressing the respective RSFP were first irradiated with light of 405 nm for varying time periods (between 0 and 300 ms), and subsequently the on-switching by light of 488 nm was recorded. From these responses, we determined the relative residual fluorescence intensity to which the fluorophores were transferred (Fig. 3b). At an intensity of 4.1 kW/cm^2^ of 405 nm light, Padron2 was switched to a residual fluorescence intensity of 5 % within 5.3 ms, while switching of Padron required nearly 12-times longer irradiation to be switched to the same value. The fluorescence of bacterial colonies expressing Kohinoor could not be switched to a residual fluorescence intensity below 8.2 % and switching times to achieve this value were two orders of magnitude longer than those required for Padron2.

As an additional benefit, the ensemble off-switching speed of Padron2 increased with higher 405 nm intensity (Suppl. Fig. 5a). This is an important feature of RSFPs for RESOLFT nanoscopy, as a predictable response of switching kinetics to increasing light intensities enhances the robustness of the imaging protocol and allows to shorten the imaging dwell time by increasing the laser intensity, if required. Increasing the 405 nm light intensity enabled us to reach residual off-state fluorescence intensities below 1 % with Padron2 (Suppl. Fig. 5a). In contrast, increasing the 405 nm laser intensity was not practical when using Padron or Kohinoor, as it resulted in increased residual fluorescence intensities (Suppl. Fig. 5b,c).

With respect to on-switching, Padron2 exhibits a fast switching half-time of 32.8 ms at 488 nm (1.3 kW/cm^2^). Thereby it is slightly faster than Kohinoor (37.6 ms) and strongly outperforms Padron (81.9 ms) (Fig. 3c; Tab.1).

Taken together, Padron2 outperforms both Padron and Kohinoor at all tested switching conditions. Most notably, it exhibits an outstanding resistance against switching fatigue as well as fast and robust switching to the off-state.

### RESOLFT nanoscopy

Most previous RESOLFT studies relied on negative switching RSFPs, the most prominent being rsEGFP2. In a beam-scanning RESOLFT approach using rsEGFP2, the fluorophores are initially switched to the on-state by a regularly focused beam of 405 nm, followed by patterned deactivation of peripheral fluorophores, which is usually facilitated with a doughnut-shaped laser beam of 488 nm. A regularly shaped laser beam of 488 nm then probes the fluorescence of RSFPs remaining in the fluorescent on-state at the center of the doughnut. With Padron2, using a sequential imaging scheme, we were able to perform RESOLFT nanoscopy on living cells using a sequence of switching steps (Fig. 4).

**Figure 4.**
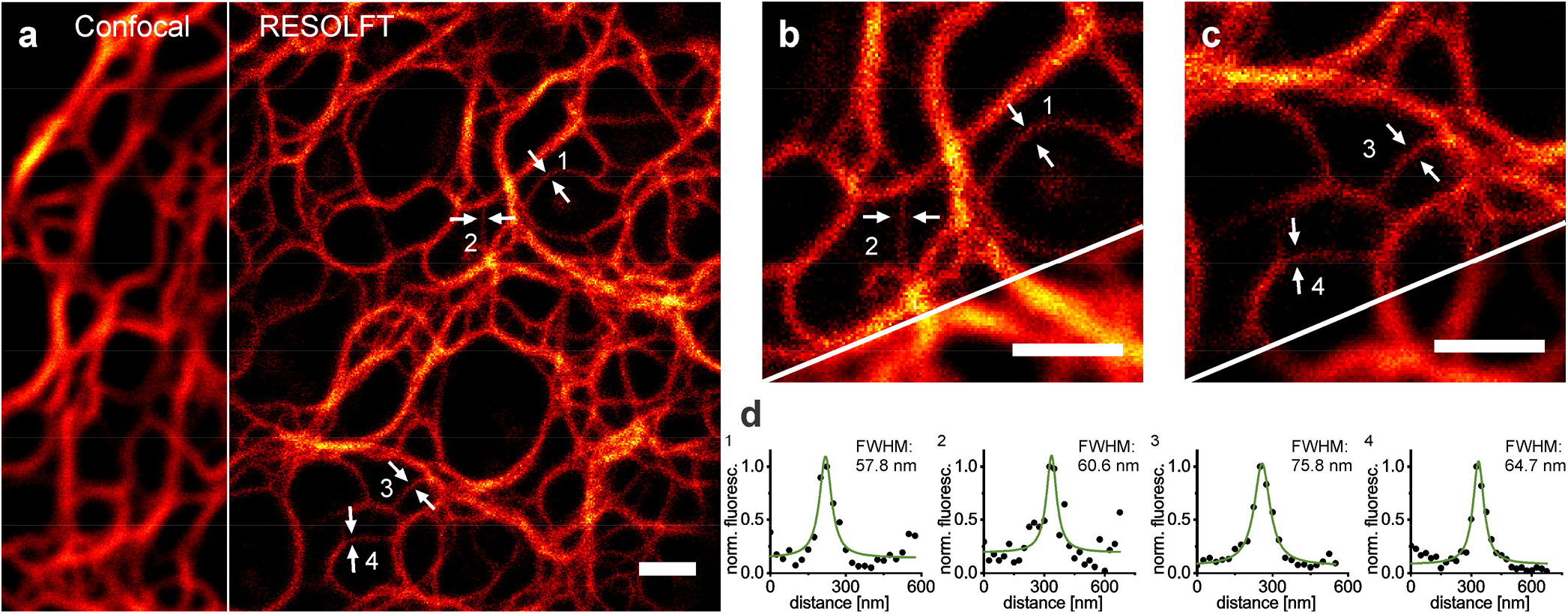
Confocal microscopy and RESOLFT nanoscopy image of vimentin-Padron2 fusion constructs. **(a)** Confocal and RESOLFT overview image. **(b, c)** Detail view of intensity line profiles 1 and 2 (b) or 3 and 4 (c). **(d)** Intensity line profiles of the positions indicated by arrows. HeLa cells were transiently transfected with the expression plasmid and were imaged at ambient temperature 24 h post transfection. The RESOLFT image was recorded with sequential irradiation with 70 μs 488 nm activation with a regularly focused beam, 350 μs off-switching with a doughnut-shaped beam, and 120 μs 488 nm readout with a regularly focused beam. Scale bars: 1 μm, 25 nm pixel size. Images are raw data. Line profiles were measured across 3 adjacent pixels and were fitted with a Lorentzian function.

Using negative switching RSFPs, it is unavoidable to use a sequence of on-switching, off-switching and fluorescence readout steps because the latter two processes are triggered by the same wavelength. However, with positive switching RSFPs, off-switching and fluorescence excitation are induced by different wavelengths. Thus, such a sequence of switching and readout steps is not necessarily required with positive switching RSFPs. We therefore aimed at establishing a live-cell RESOLFT imaging protocol which entirely refrains from sequential illumination steps, but rather uses a regularly focused excitation beam of 488 nm that is superimposed with a doughnut-shaped beam of 405 nm light. In this concept, RSFPs in the periphery of the focus predominantly are present in the off-state because in the doughnut-shaped focus off-switching dominates over the on-switching process. Towards the central focal region, where the 405 nm light intensity is at a minimum, the intensity of light of 488 nm increases. Hence, towards the center of the doughnut, increasingly more RSFPs reside in the on-state and emit fluorescence. By tuning the relative light intensities of 405 and 488 nm the size of the central, fluorescent region can be adjusted to a sub-diffraction sized volume. Because this approach requires no sequential irradiation steps, we refer to this imaging concept as one-step RESOLFT nanoscopy.

This approach to RESOLFT nanoscopy requires a positive switching RSFP, which, if simultaneously irradiated with light of 405 nm and 488 nm, is mostly present in the off-state. Presumably, when irradiated with both wavelengths the positive switching RSFP will cycle repeatedly between the on- and the off-state. Hence, a suitable RSFP is required to display a good resistance against switching fatigue. We anticipated that Padron2 fulfills these requirements. To further investigate the suitability of Padron2 for this approach, we irradiated bacterial colonies expressing Padron2, Padron, or Kohinoor with light of both 405 nm and 488 nm simultaneously. Intensities were chosen so that the off-switching process was dominant and the proteins were effectively switched off, a situation that would be required for one-step RESOLFT nanoscopy. In this experiment, Padron2 displayed faster off-switching rates as well as lower residual fluorescence intensities than both Padron and Kohinoor (Suppl. Fig. 6). In addition, photodestruction of Padron2 was very low in comparison to Padron and Kohinoor when illuminated with both wavelengths at the same time (Suppl. Fig. 7). These observations suggested that Padron2, but neither Padron nor Kohinoor, would be suitable for RESOLFT nanoscopy without sequential irradiation steps.

To evaluate this RESOLFT implementation on living cells with Padron2, we expressed the RSFP fused to the intermediate filament protein vimentin (VIM-Padron2) or the nuclear-pore protein Nup50 (Padron2-NUP50) in human HeLa cells (Fig. 5). The resulting images revealed details that were concealed in the corresponding diffraction-limited confocal images. On vimentin filaments we typically measured a full width at half maximum (FWHM) of ~75 nm and we were able to resolve structures that were as close as ~140 nm. Padron2-Nup50 was expressed to a much lower level than VIM-Padron2. To be able to image these dim samples, we made use of the fact that with superimposed irradiation of doughnut and regularly focused fluorescence excitation the fluorophores in the periphery of the doughnut region are mostly present in the off-state, while central ones remain fluorescent. Hence, the pixel dwell-time can be prolonged to collect more photons if required. We utilized this property and prolonged the pixel dwell time from 300 μs to 500 μs. Also, the images of the nuclear pores clearly demonstrated that with this simplified RESOLFT approach the diffraction barrier can be overcome in living cells.

**Figure 5.**
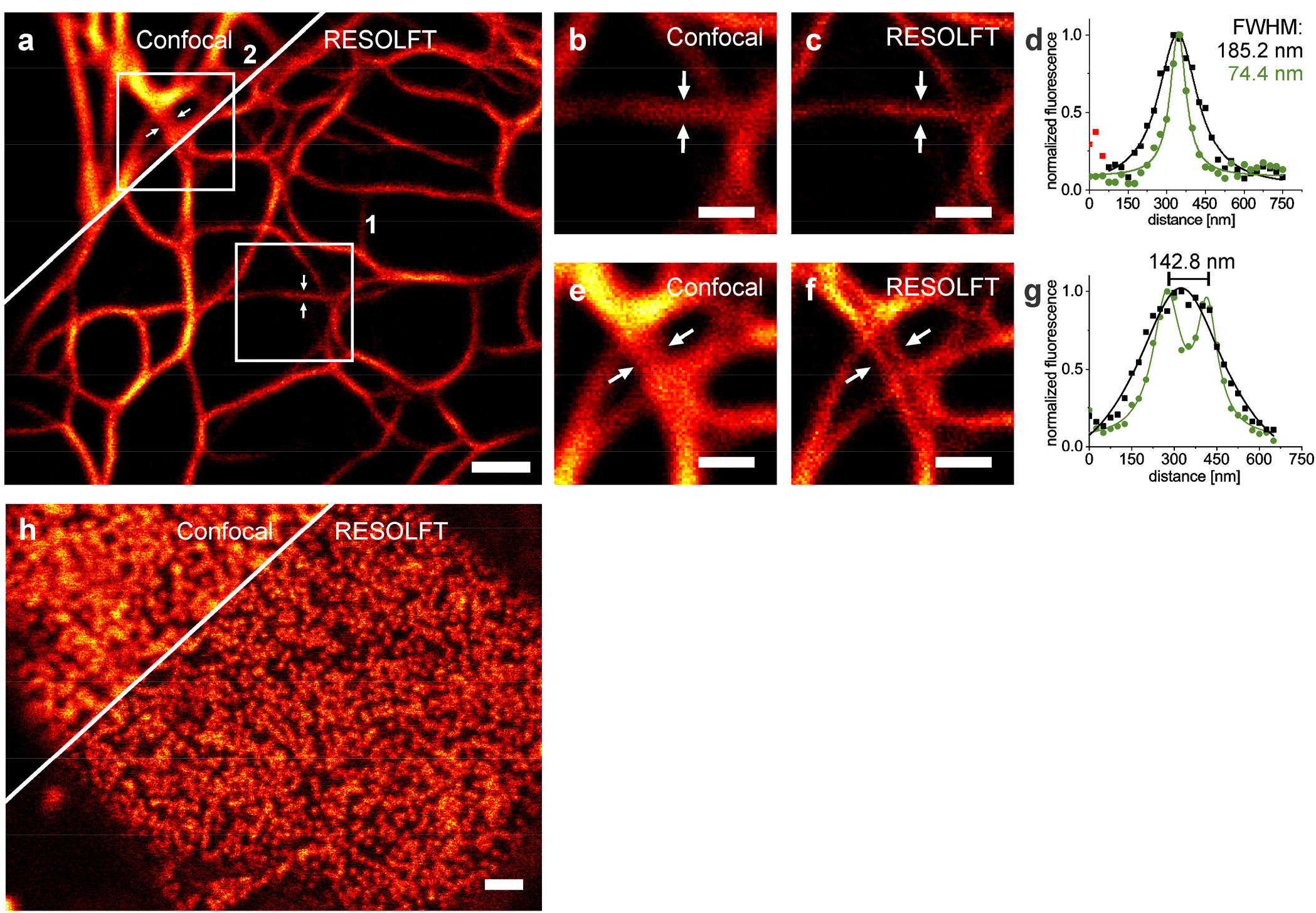
Confocal microscopy and one-step RESOLFT nanoscopy images of Padron2 fusion constructs. **(a)** Confocal and RESOLFT overview image of vimentin-Padron2. **(b)** Confocal and **(c)** RESOLFT image with **(d)** intensity line profile measured across 3 adjacent pixels at the position marked by the arrows of boxed region 1 in (a). **(e)** Confocal and **(f)** RESOLFT image with **(g)** intensity line profile measured across 3 adjacent pixels at the position marked by the arrows of boxed region 2 in (a). **(h)** Confocal and RESOLFT image of Padron2-Nup50. HeLa cells were transiently transfected with the respective expression plasmid and imaged at ambient temperature 24 h post transfection. The RESOLFT image in (a) was recorded with a pixel dwell time of 300 μs with constant superimposed irradiation of a regularly focused 488 nm laser at 1.4 kW/cm^2^ and a doughnutshaped 405 nm laser at 1.0 kW/cm^2^. Peaks were fitted with a Lorentzian function. The RESOLFT image in (h) was recorded with a pixel dwell time of 500 μs with constant superimposed irradiation of a regularly focused 488 nm laser at 1.7 kW/cm^2^ and a doughnut-shaped 405 nm laser at 0.9 kW/cm^2^. Scale bars: 1 μm (a, b, h), 0.5 μm (c, d, f, g). Pixel size: 25 nm. All images are raw data.

## Discussion

In this study, we engineered the new positive switching RSFP Padron2 and achieved improvements with regard to properties that were identified as rather unfavorable in established green positive switching RSFPs, while maintaining the beneficial properties of its template Padron. Padron2 specifically stands out with respect to switching fatigue, resistance against photobleaching in the on-state and it exhibits fast ensemble switching. In the ensemble, it can be reversibly switched to very low residual fluorescence intensities. The molecular brightness of Padron2 has been reduced by a factor of 3.2 or 1.2 compared to Padron or Kohinoor, respectively, but this is outweighed by its improved expression at 37 °C, a key requirement for the use of a genetically encoded probe in mammalian cells.

Previously, negative switching RSFPs have outperformed the available positive switching RSFPs with respect to most relevant characteristics. The switching performance of Padron2 is similar to that of rsEGFP2,^11^ the most widely used negative switching RSFP for RESOLFT nanoscopy. Padron2 sustains half of the switching cycles reported for rsEGFP2. Its molecular brightness is lower by about a factor of 2.3. The ensemble switching speed of both RSFPs is comparable, and with Padron2 a much lower residual off-state fluorescence intensity could be reached than reported for rsEGFP2.

Padron2 further stands out by the fact that it is photostable when irradiated simultaneously with light of both its off- and on-switching wavelength (405 and 488 nm, respectively). While Padron and Kohinoor exhibited high switching fatigue when irradiated with both wavelengths simultaneously, Padron2 fluorophores could undergo many switching cycles under these conditions. When Padron2 is irradiated with suitable intensities of both wavelengths, the vast majority of the molecules are in the off-state; when the ratio of intensities is shifted towards 488 nm, increasingly more Padron2 proteins are present in the on-state and emit fluorescence. This switching behavior was exploited to implement a RESOLFT scheme without sequential switching.

One-step RESOLFT nanoscopy is thereby similar to STED nanoscopy,^1^ which also does not require sequential irradiation steps. STED nanoscopy, however, requires light intensities which are up to 6 orders of magnitude higher than typically used in RESOLFT nanoscopy.^34^ The first demonstration of RESOLFT nanoscopy in 2005 used the one-step RESOLFT concept in order to achieve diffraction-unlimited resolution.^7^ Since then, this approach had not been explored further because of a lack of positive switching RSFPs suitable as fusion tags for live cell imaging. Instead, RESOLFT nanoscopy was established with negative switching RSFPs relying on sequential irradiation schemes as a seemingly inevitable compromise.

Conceptually, one-step RESOLFT nanoscopy with positive switching RSFPs has distinct advantages, namely the simplicity of the recording scheme and the fact that the number of detected photons can be increased by prolonging the pixel dwell time because the superposition of light patterns maintains the state separation of fluorophores in the on- and off-state.

With Padron2, we were able to perform one-step RESOLFT nanoscopy in living cells. The implementation does not require data processing since the detected photons are directly mapped to a pixel in the image. The recoding times are comparable to those reported with rsEGFP2, but were realized with lower overall light doses. The resolution obtained is not as good as typically achieved with rsEGFP2 in sequential RESOLFT. Possibly, this lack of resolution attained resulted from challenges associated with the generation of a small zero due to the competitive switching close to the zero.

In summary, with Padron2 we have generated a positive switching RSFP, which outperforms all previous RSFPs of this class. Padron2 allows the implementation of live-cell RESOLFT nanoscopy without the need for sequential irradiation steps.

## Methods

### Mutagenesis

Coding sequences were mutagenized by site-directed saturation (QuickChange™ protocol, Stratagene, La Julla, Ca, USA), error-prone35 or multiple-site mutagenesis.36 GFPends were introduced via PCR and re-insertion of the coding sequence into the backbone via restriction with enzymes EcoRI and XhoI (forward primer: ACGGATCCAATGGTGAGCAAGGGCGAGGAGAACAACATGGCCGTGATTAAACCAGAC, reverse primer: ATTAAGCTTCGAATTCTTACTTGTACAGCTCGTCCATGGCCTGCCTCGGCAG).

### Screening

Mutant libraries of early variants were generated in the pQE-31 expression vector (Qiagen, Hilden, Germany) and expressed in SURE *E. coli* (Stratagene, CA, USA) on LB-agar. Mutant libraries of later variants were generated in the pBAD/HisB expression vector from pBAD-mKalama1, which was a gift from Robert Campbell (Addgene plasmid #14892).^37^ Libraries were transformed into One Shot TOP10 Electrocomp *E.coli* (Thermo Fisher Scientific, Waltham, MA, USA) and plated on LB agar with 0.02 % (w/v) arabinose. Bacterial colonies were screened with a customized DM5500B microscope (Leica Microsystems, Wetzlar, Germany) with a SCAN 100×100 stage (Märzhäuser Wetzlar, Wetzlar, Germany). Laser sequences were controlled with an FPGA program based on LabVIEW (National Instruments, Austin, TX).

### Protein purification

Padron, Padron2, and Kohinoor were expressed in One Shot TOP10 Electrocomp *E.coli* (Thermo Fisher Scientific, Waltham, MA, USA) from the pBAD/HisB expression system. For this, the coding sequence for Kohinoor was PCR amplified from Kohinoor/pRSETb (forward primer: AGGGCTCGAGCATGAGTGTGATTAAACC, reverse primer: AACGAATTCTTACTTGGCCTGCCT), which was a gift from Takeharu Nagai (Addgene plasmid #67770),^24^ and inserted into pBAD/HisB with restriction enzymes EcoRI and XhoI. After 24 h at 37 °C (Padron2, Kohinoor) or 48 h at 30 °C (Padron) growth on LB-agar with 0.02 % (w/v) arabinose, cultures were kept at room temperature over night to ensure full maturation. Cells were collected in 20 mM phosphate, 500 mM NaCl and 20 mM imidazole (pH 7.4). Cell suspensions were incubated on ice for 4 h with 1 mg/ml lysozyme (Serva electrophoresis, Heidelberg, Germany). After incubation, complete protease inhibitor (Roche, Basel, Switzerland) was added and samples were frozen and thawed 5x in liquid nitrogen and lukewarm water. Lysates were then centrifuged with 0.5 μl benzonase (Thermo Fisher Scientific, Waltham, MA, USA) added per 2 ml at 4 °C and 21,000 rcf for 3–6 h. RSFPs were isolated from the supernatant with the His SpinTrap kit (GE Healthcare, Chicago, IL, USA) and concentrated with Vivaspin 500 centrifugal concentrators with a molecular weight cut-off of 10,000 kDa (Sartorius, Göttingen, Germany). Elution buffer was exchanged to standard Tris protein buffer (100 mM Tris and 150 mM NaCl at pH 7.5) with NAP-5 columns (GE Healthcare, Chicago, IL, USA) and samples were concentrated as before.

### Size exclusion chromatography

Protein samples were diluted to 10 μM and equilibrated at 6 °C overnight. 250 μl were applied to an Äkta pure chromatography system equipped with a Superdex 200 Increase 10/300 GL column (GE Healthcare, Chicago, IL, USA) and run at 6 °C. Flow rate was 0.75 ml/min and protein elution was monitored with the U9-L UV monitor (GE Healthcare, Chicago, IL, USA) at 280 nm.

### Spectroscopy

Purified protein samples were diluted to an absorption of 0.1 at 280 nm, corresponding to a concentration of approximately 20 μM. Samples were equilibrated over night at 21 °C. Absorption spectra were recorded with a Cary 4000 UV-VIS spectrophotometer (Varian, Palo Alto, CA, USA) in an ultra-micro fluorescence cell cuvette with a 1.5 mm light path (Hellma, Müllheim, Germany). Spectra were normalized to absorption at 280 nm and smoothed in OriginPro 2018b (OriginLab Corporation, Northhampton, MA, USA) with a Savitzky-Golay filter with 10 point window size. Emission spectra of switching states (Suppl. Fig. 3d-f) were measured in the same cuvette with a Cary Eclipse fluorescence spectrophotometer (Varian, Palo Alto, CA, USA) with excitation at 460 nm. Switching to the on- or the off-state was facilitated in a cuvette with a mercury-vapor lamp with a HQ405/10 X filter for off-switching (9.9 mW/cm^2^, 0.74 mW measured behind the cuvette filled with Tris protein buffer) and a ET500/20 X filter for on-switching (18.7 mW/cm^2^, 1.4 mW measured behind the cuvette filled with Tris protein buffer). Proteins were switched for 3–5 min until saturation. Power was measured with a PM200 powermeter equipped with a S170C sensor (ThorLabs, Newton, NJ, USA). The spectra recorded this way were integrated and used for the determination of the residual off-state and equilibrium state fluorescence intensity in solution.

The extinction coefficient was calculated for the on-state of RSFPs from comparative measurements with mEGFP. On-state absorption of the deprotonated peak in normalized absorption spectra was used following eq. 1.1 with an extinction coefficient of 56,000 M^−1^cm^−1^ for mEGFP.^38^ Due to the influence of the amino acid composition of proteins during normalization a correction term was added to the equation to correct for differing absorption of aromatic amino acids, which was calculated with the ExPASy ProtParam online tool. 6–7 absorption spectra were used and calculated extinction coefficients were averaged.

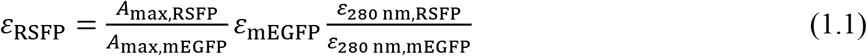

(ε, extinction coefficient, A, absorption)

Quantum yield was calculated from the integrated on-state emission spectra relative to the quantum yield of Padron following equation 1.2. 3–4 spectra were used for the calculation and results were averaged.

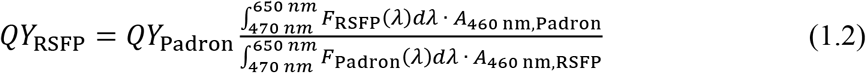

(QY, quantum yield; F(λ), fluorescence emitted with the wavelength λ; A, absorption)

The excitation and emission spectra in Fig. 1a and Suppl. Fig. 3e,f and pH spectra were recorded in 96 well UV-Star micro plates (Greiner Bio-One, Frickenhausen, Germany) with the Cytation 3 imaging plate reader (BioTek, Winooski, VT, USA). Samples were diluted to 200 μM, equilibrated at 21 °C over night and diluted to 200 μl 5 μM in triplicate wells prior to measurements. Several buffers in 0.5 pH steps were used for dilution (pH 3.0–5.5: 100 mM citric acid, 150 mM NaCl; pH 6.0–7.0: 100 mM KH_2_PO_4_, 150 mM NaCl; pH 7.5–8.5: 100 mM Tris, 150 mM NaCl; pH 9.0–10.5 (+10.25): 100 mM glycine, 150 mM NaCl). Emission spectra were recorded with 470 nm excitation from 500 to 700 nm, excitation spectra from 400 to 520 nm with emission detection at 550 nm. Detector sensitivity was scaled to the brightest well. For p*K_a_* calculation, fluorescence was recorded with at 485/20 excitation filter and a 528/20 filter. Fluorescent pH response was fitted to a mono-(Padron; eq. 1.3) or biphasic (Padron2, Kohinoor; eq. 1.4) dose response function.

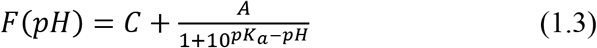

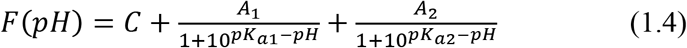

(*F*, pH-dependent fluorescence intensity, *C* and *A* are fitted parameters.)

### Fluorescence lifetime

Fluorescence lifetimes were measured with a Quantaurus-Tau fluorescence lifetime spectrometer (Hamamatsu, Hamamatsu, Japan). Measurements were performed in quartz-cuvettes with 470 nm excitation and detection at 516 nm. 10,000 photons were recorded after measurement of the internal response function with Polybead amino 0.10 micron microspheres (Polysciences, Warrington, PA, USA). Data was analyzed with version 3.0.0.80 of the Quantaurus-Tau software.

### Switching characterization

Switching kinetics and associated parameters were measured with the automated screening microscope in bacterial colonies. To ensure a maximal fluorescence signal, an autofocus was employed. Photobleaching was recorded with continuous 488 nm irradiation at 2.3 ± 0.3 kW/cm^2^ with 3–4 averaged repetitions with 5–10 colonies each.

Residual off-state fluorescence intensity was quantified by switching proteins off with progressively increasing 405 nm irradiation duration at different intensities with 5 colonies per condition in 2–3 repetitions. Residual off-state fluorescence intensities were calculated from the consecutive on-switching curve. Graphs display averaged data with an ExpDec2 nonlinear curve fit calculated in OriginPro 2018b (OriginLab Corporation, Northampton, MA, USA). Off-switching intensities are indicated in the respective graphs, on-switching was performed at 20.2 ± 0.9 kW/cm^2^. On-switching curves shown in Fig. 3c were analyzed by switching proteins to 5 % (Padron, Padron2) or 10 % (Kohinoor) residual off-state fluorescence intensity with 405 nm and subsequent activation with 488 nm irradiation at 1.3 kW/cm^2^. Three repetitions were measured with 10 colonies each. Measurements were averaged and normalized.

Switching fatigue was determined by repeatedly cycling the proteins between the on- and off-state using light doses that switched the proteins off to 5 % (Padron, Padron2) or 10 % (Kohinoor) residual fluorescence intensity and on to 95 % of their maximal on-state fluorescence intensity. Off-switching 405 nm light intensity was 3.6 kW/cm^2^, 488 nm intensity was 2.6 kW/cm^2^; 3 repetitions were measured with 10 colonies each. End points of the activation curves were used for illustration of the switching fatigue after normalization and averaging. As all three RSFPs differed with regards to switching speed, light doses varied in switching fatigue measurements. Irradiation duration and light doses are listed in Suppl. Tab. 1. To estimate the influence of differing light doses, we repeated the measurements for Padron2 with Kohinoor irradiation settings and vice versa (Suppl. Fig. 8).

Switching RSFPs off using simultaneous irradiation was recorded as follows: Proteins were subjected to a single full switching cycles with sequential irradiation (10 ms 405 nm irradiation at 4.1 ± 0.1 kW/cm^2^, 50 ms 488 nm irradiation at 17.4 ± 0.3 kW/cm^2^). Subsequently, 10 cycles of switching to the off-state with simultaneous 405 and 488 nm irradiation for 100 ms and switching to the on-state with 488 nm alone for 50 ms were recorded. For off-switching, different intensity combinations were measured (405 nm at 4.1 ± 0.1 kW/cm^2^ (low) or 56.2 ± 1.2 kW/cm^2^ (high), 488 nm at 1.1 ± 0.1 kW/cm^2^ (low) or 17.4 ± 0.3 kW/cm^2^ (high)), 488 nm irradiation for the on-switching was at 17.4 ± 0.3 kW/cm^2^ in all measurements. For every intensity combination, 3 repetitions were measured with 10 colonies each. Off-switching curves shown in supplemental figure 6 are averaged curves, graphs with 10 consecutive cycles in supplemental figure 7 are representative single measurements.

For the evaluation of protein expression after growth at different temperatures, fluorescence intensities were probed with a 488 nm excitation intensity of 18.6 ± 0.5 kW/cm^2^ after growth at 37 °C for 24 h or 30 °C for 48 h.

Laser powers used were measured behind the objective lens (N PLAN L 20s/0.40, Leica Microsystems, Wetzlar, Germany) with a LabMax-TO laser power meter equipped with an LM-2 VIS sensor (Coherent, Santa Clara, CA, USA). For calculation of laser intensities, focal spot sizes were measured by focusing on a mirror surface on the microscope stage. Reflected light reached the detection path by usage of a 50/50 beam splitter, where a mirror was inserted and the spot was visualized with a SPC900NC webcam sensor (Philips, Amsterdam, Netherlands). Pixel size was calibrated with the image of a 10 μM scale, and intensity line profiles of the focal laser spots were measured. FWHM values were calculated and a circular area with the FWHM as diameter was assumed for intensity calculations.

### Cloning

Mammalian expression vectors were created by cloning after PCR amplification of the Padron2 coding sequence with primers listed in Suppl. Tab. 2.

pVimentin-Padron2: The coding sequence of Padron2 was inserted into pmKate2-vimentin (Evrogen, Moscow, Russia) with restriction enzymes AgeI and NotI, replacing mKate2. pKeratin-Padron2: The RSFP coding sequence of Padron2 was inserted into pTagRFP-keratin (Evrogen, Moscow, Russia) with restriction enzymes KpnI and NotI, replacing TagRFP.

Actin-filaments were labelled indirectly with Padron2 fused to Lifeact. The coding sequence was inserted into lifeact-EGFP pcDNA3.1(+)^39^ with enzymes BamHI and NotI, replacing EGFP to create pLifeact-Padron2. Microtubules were indirectly labelled with a fusion construct of Map2 and Padron2 in pPadron2-Map2. For this, the coding sequence for Map2 was inserted into the backbone of pEGFP-Tub (BD Biosciences Clontech, Franklin Lakes, NJ, USA) with restriction enzymes XhoI and BamHI after amplification from pDONR223-MAP2^40^ (forward primer: GATCTCGAGTGATGGCAGATGAACGGAAAGACGAAGC, reverse primer: GGTGGATCCTTATCACAAGCCCTGCTTAGCGAGTGCAGC), replacing the tubulin sequence. The mEGFP sequence was subsequently replaced by the one for Padron2 with NheI and BglII.

For expression in the endoplasmic reticulum, the Padron2 coding sequence was inserted into *pEF/myc/ER* (Invitrogen Life Technologies, Carlsbad, CA, USA) with enzymes SalI and NotI to create pPadron2-ER, which added an N-terminal ER signal peptide and a C-terminal ER retention signal to the RSFP. Cytosolic expression was facilitated by expression of free Padron2 inserted into the TagRFP-N vector (Evrogen, Moscow, Russia) with enzymes AgeI and NotI, replacing TagRFP to create pPadron2-N. Labelling of the nuclear pore complex was done with the expression plasmid pPadron2-Nup50 coding for a fusion construct. It was created by replacing the sequence of mEmerald with that of Padron2 with enzymes NheI and BglII in mEmerald-Nup50-C-10, which was a gift from Michael Davidson (Addgene plasmid #54209). Histones were labelled by fusing Padron2 to H2bn in pPadron2-Histon1 (H2bn). The EGFP sequence in pEGFP-Hist1H2BN^41^ was replaced with enzymes NheI and BglII. For Mitochondrial localization, the DsRed coding sequence in pDsRed1-Mito (BD Biosciences Clontech, Franklin Lakes, NJ, USA) was replaced with that of Padron2 with enzymes AgeI and NotI creating pMito-Padron2. Centromeres were targeted with centromere protein C (CenpC) fusion constructs from plasmid pPadron2-CenpC1. The coding sequence for the target protein was PCR amplified from pDONR223-CenpC^40^ (forward primer: CAGATCTCGAGTGGCTGCGTCCGGTCTGGA, reverse primer: TCCGGTGGATCCTTAGCATTTCAGGCAACTCTCCT) and inserted into pEGFP-Tub (BD Biosciences Clontech, Franklin Lakes, NJ, USA) with enzymes XhoI and BamHI replacing the tubulin sequence. The EGFP sequence was subsequently replaced with that of Padron2 with enzymes NheI and XhoI. Targeting of caveolae was facilitated by inserting the coding sequence for caveolin-1 after PCR amplification from pDONR223-CAV1^40^ (forward primer: TCCGCTAGCATGTCTGGGGGCAAAT, reverse primer: CCGGTGGATCCCGGGCCCGCGGTATTTCTTTCTGCAAGTTGATG) into the pPadron2-N with enzymes NheI and BamHI creating pCaveolin1-Padron2. For peroxisomal targeting, Padron2 was inserted into pEGFP-PTS^41^ with enzymes NheI and BglII, replacing the EGFP sequence to create pPadron2-peroxi. This expression plasmid adds a C-terminal peroxisomal targeting sequence (PTS) to Padron2.

### Microscopy

All images shown were recorded with transiently transfected HeLa cells 24 h post transfection. Cells were mounted in DMEM without phenol red (Thermo Fisher Scientific, Waltham, MA, USA) and imaged at ambient temperature with a customized 1C RESOLFT QUAD scanning microscope (Abberior Instruments, Göttingen, Germany). The microscope was equipped with a UPLSAPO 1.4 NA 100x oil immersion objective (Olympus, Shinjuku, Japan) as well as a 405 nm and a 488 nm continuous-wave laser (both Cobolt, Solna, Sweden). The 405 nm doughnut-shaped beam was realized with an easy 3D module (Abberior Instruments, Göttingen, Germany). Fluorescence was detected with a SPCM-AQRH-13 photon counting module (Excelitas Technologies, Waltham, MA, USA) with a HC 550/88 detection filter. Laser powers were measured behind the objective with a PM200 power meter with the S170C sensor (ThorLabs, Newton, NJ, USA). The circular or ring-like area of both beams at FWHM intensity in the focus were determined and used for further calculations. Images and filament intensity line profiles measured with 3 adjacent lines were analyzed with the Fiji distribution of imageJ (v1.52p)^42, 43^ and OriginPro 2018b (OriginLab Corporation, Northhampton, MA, USA).

## Acknowledgements

We thank S. Schnorrenberg for helpful discussions and S. Löbermann and T. Gilat for excellent technical support.

## Supporting information

Supplemental figures and tables.

## Abbreviations

RESOLFT: reversible saturable optical linear (fluorescence) transitions
RSFP: reversibly switchable fluorescent proteins
STED: stimulated emission depletion

