## Supplemental Information for "The Positive Switching RSFP Padron2 Enables Live-Cell RESOLFT Nanoscopy Without Sequential Irradiation Steps"

```
                •••••••••1••••••••10••••••••20••••••••30••••••••40••••••••50••••••••60••••••••70
Padron          ---------MSVIKPDMKIKLRMEGAVNGHPFAIEGVGLGKPFEGKQSMDLKVKEGGPLPFAYDILTMAFCYGNRVFAKY
Kohinoor        ---------MSVIKPDMKIKLRMEGAVNGHPFAIEGVGLGKPFEGKQSMDLKVKEGGPLPFAYDILTMAFCYGNRVFAKY
Padron2         MVSKGEENNMAVIKPDMKIKLRMEGAVNGHPFAIEGVGLGKPFEGKQSVDLKVKEGGPLPFAYDILSMAFCYGNKVFIKY

                ••••••••80••••••••90••••••••100•••••••110•••••••120•••••••130•••••••140•••••••150
Padron          PENIVDYFKQSFPEGYSWERSMIYEDGGICNATNDITLDGDCYIYEIRFDGVNFPANGPVMQKRTVKWELSTEKLYVRDG
Kohinoor        PENIVDYFKQSFPEGYSWERSMIYEDGGICIATNDITLDGDCYIYEIRFDGVNFPANGPVMQKRTVKWEPSTEKLYVRDG
Padron2         PENIVDYFKQLFPEGYSWERSMIYEDGGICNATNDITLDGDCFIYEIRFDGVNFPANGPVMQKRTVKWEPSTEKLYVRDG

                ••••••••160•••••••170•••••••180•••••••190•••••••200•••••••210•••••••220•••••••
Padron          VLKSDGNYALSLEGGGHYRCDFKTTYKAKKVVQLPDYHSVDHHIEIKSHDKDYSNVNLHEHAEAHSELPRQAK-----
Kohinoor        VLKSDGNYALSLEGGGHYRCDSKTTYKAKKVVQLPDYHDVVHHIEIKSHDRDYSNVNLHEHAEAHSGLPRQAK-----
Padron2         VLKSDGNYALSLEGGGHYRCDSKTTYKAKKVVQLPDYHAVDHHIEIKSHDKDYSNVNLHEHAEAHSGLPGQAMDELYK
```

**Supplemental Figure 1.** Protein sequence alignment of Padron, Padron2, and Kohinoor. Amino acids are numbered in accordance with the sequence of Dronpa. Mutations that were introduced to Dronpa to create the original Padron are highlighted in green, mutations of Padron2 and Kohinoor in comparison to Padron are highlighted in yellow.

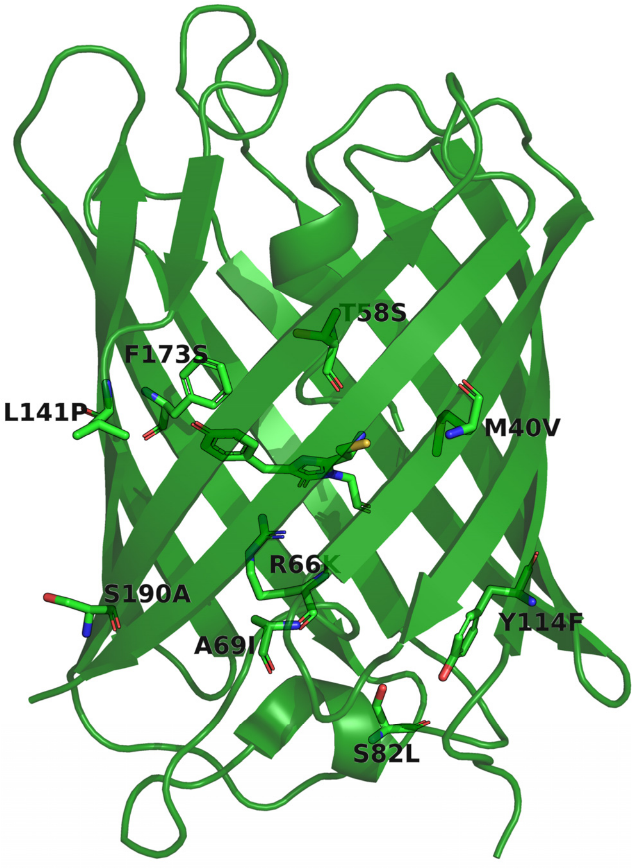


**Supplemental Figure 2.** Mutated positions in Padron. Amino acids targeted by mutagenesis during the generation of Padron2 are displayed in stick representation within the on-state structure of Padron (PDB ID: 3ZUJ), labels are mutations introduced in Padron2. R221 and E218 were not part of the solved crystal structure and are not highlighted. The structure was rendered with the PyMOL Molecular Graphics System, Version 2.3 Schrödinger, LLC.

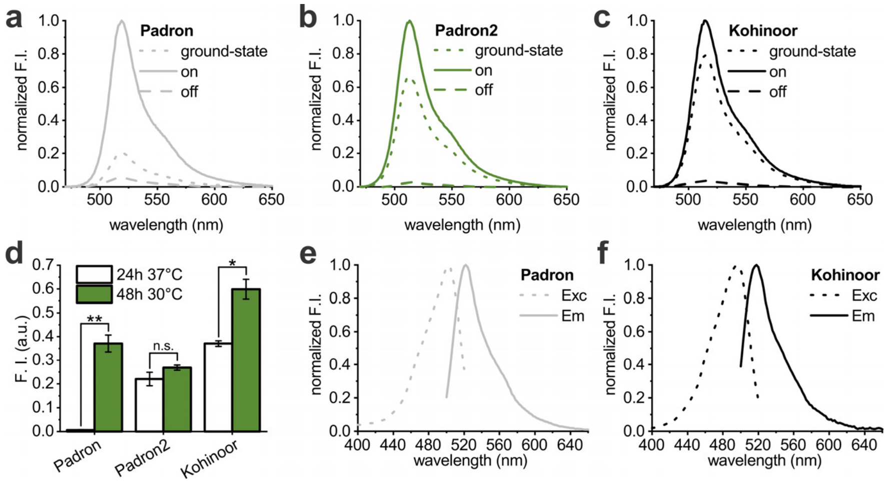


**Supplemental Figure 3.** Protein characteristics. (a, b, c) Normalized fluorescence emission spectra of the on- and off-state as well as the equilibrated state for Padron (a), Padron2 (b), and Kohinoor (c). Excitation wavelength was 460 nm. (d) Absolute fluorescence intensity values in bacterial colonies used for Fig. 1g. Padron, Padron2, and Kohinoor fluorescence intensities were probed after growth at 37 °C for 24 h and growth at 30 °C for 48 h. p-values: ** = 2.2E-6, n.s. = 0.03, * = 9.8E-5. Two sample T-Tests calculated in OriginPro 2018b (OriginLab Corporation, Northhampton, MA, USA), equal variance assumed. (e, f) Normalized Excitation (Exc) and emission (Em) spectra for Padron (e) and Kohinoor (f).

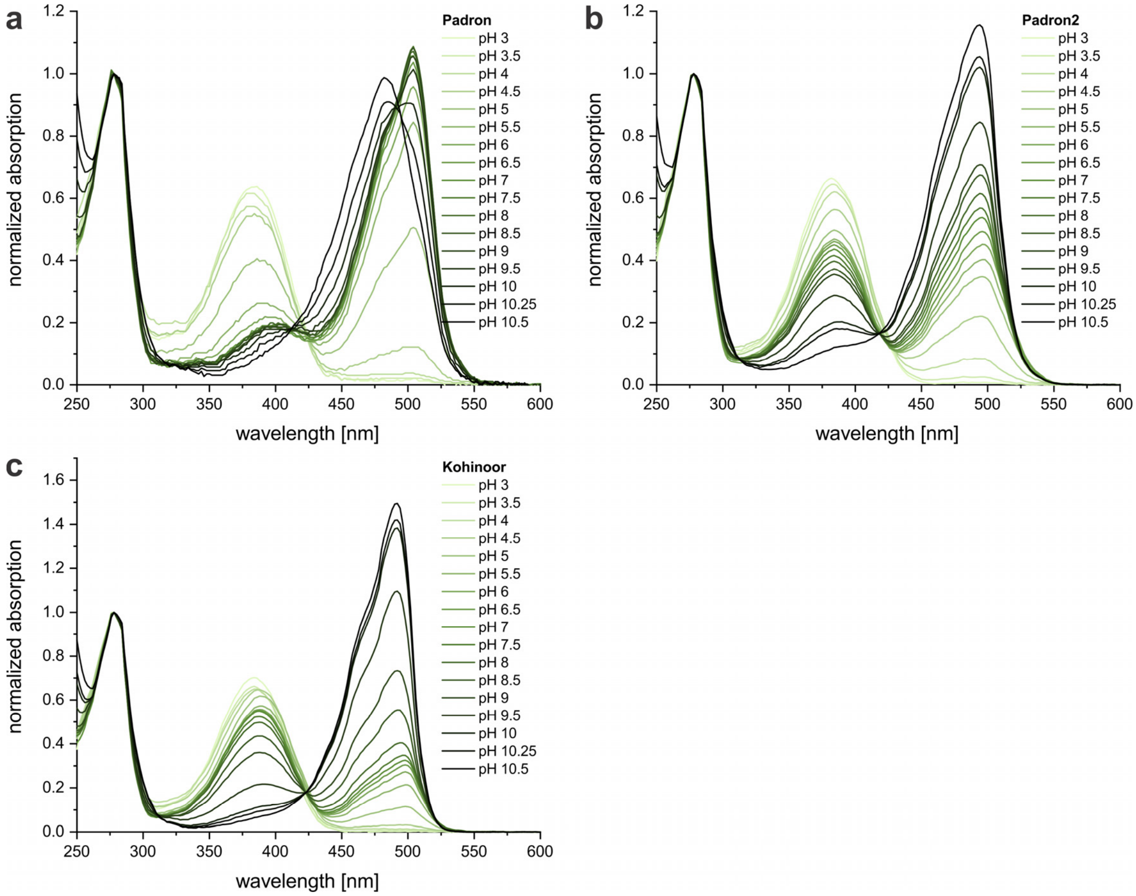


**Supplementary Figure 4.** Normalized pH absorption spectra of (a) Padron, (b) Padron2, and (c) Kohinoor. Absorption spectra were measured at the pH-values indicated in the figure legends with a plate reader in the equilibrated state at room temperature and normalized to the absorption of aromatic side chains at 278 nm.

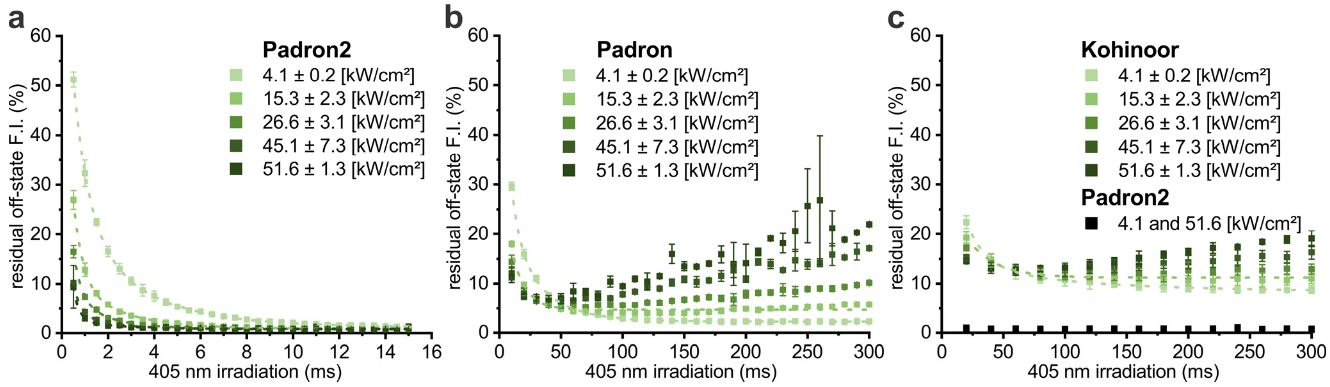


**Supplemental Figure 5.** Off-switching performance of (a) Padron2, (b) Padron, and (c) Kohinoor. Data points are residual fluorescence intensities after different 405 nm irradiation times (horizontal axes) at different intensities. Dashed lines represent exponential decay fitting data. Black squares in (c) are Padron2 data measured with the same irradiation times used for Kohinoor for two different intensities, which resulted in similar residual fluorescence intensities (data points are shown for both intensities but are overlapping).

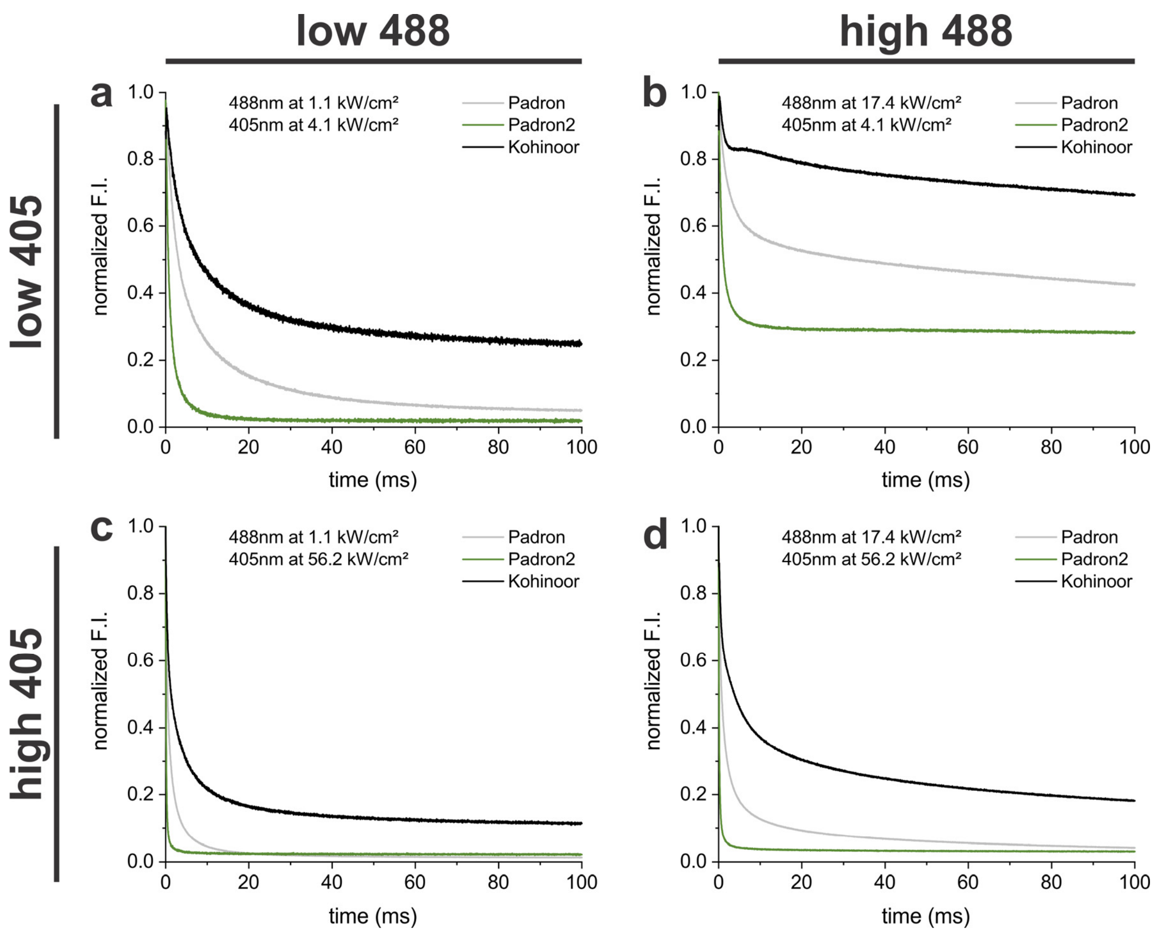


**Supplemental Figure 6.** Off-switching kinetics with simultaneous 405 and 488 nm irradiation. Fluorescence decay in bacterial colonies was measured at different intensity combinations of 405 (4.1 ± 0.1 kW/cm² (low) or 56.2 ± 1.2 kW/cm² (high)) and 488 nm (1.1 ± 0.1 kW/cm² (low) or 17.4 ± 0.3 kW/cm² (high)). Switching curves were normalized and are averaged data.

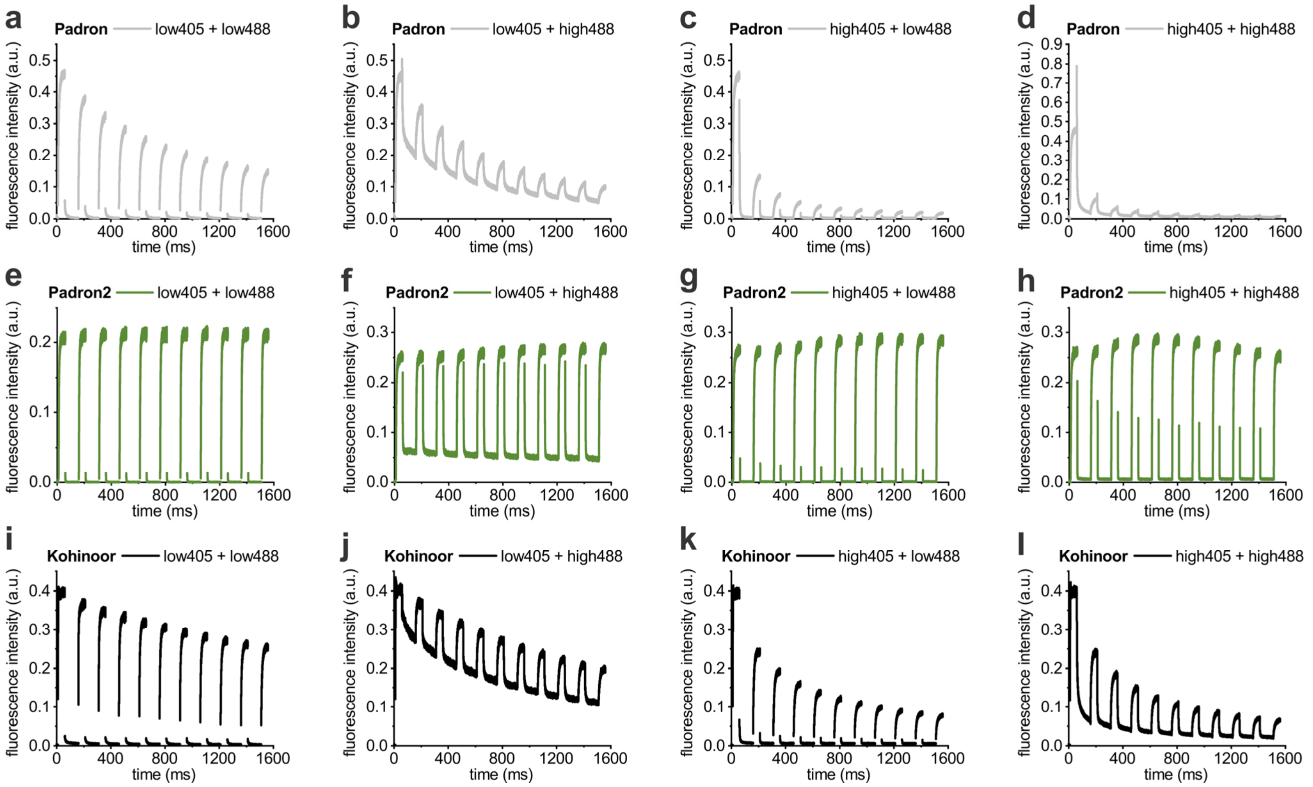


**Supplemental Figure 7.** 10 consecutive switching cycles with off-switching under simultaneous irradiation of 405 and 488 nm. Proteins expressed in bacterial colonies were initially switched off- and on again with 405 and 488 nm alone (first activation curve in graphs) and then switched to the off state for 100 ms with 405 and 488 nm at different intensity combinations (405nm: 4.1 ± 0.1 kW/cm² (low) or 56.2 ± 1.2 kW/cm² (high); 488 nm: 1.1 ± 0.1 kW/cm² (low) or 17.4 ± 0.3 kW/cm² (high)). Subsequent switching back to the on-state for 50 ms was done with the same intensity of 488 nm alone in all measurements (17.4 ± 0.3 kW/cm²). (a-d) Padron, (e-h) Padron2, and (i-l) Kohinoor. Graphs display representative measurements in a single colony.

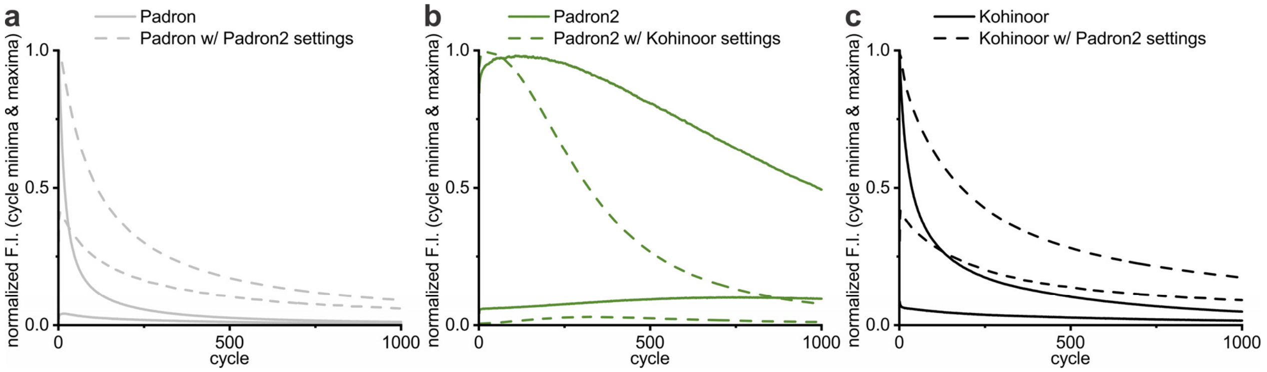


**Supplemental Figure 8.** Switching fatigue for (a) Padron, (b) Padron2, and (c) Kohinoor. Graphs include switching fatigue as shown in figure 3c (upper solid lines) with additional residual fluorescence intensity values in the off-state (lower solid lines). Dotted lines represent equivalent data of on-state and off-state fluorescence intensities after switching with the light doses applied for Padron2 and Kohinoor, respectively (cf. figure legends, Suppl. Tab. 1).

**Supplemental Table 1.** Light doses applied for switching fatigue measurements.

| | Irradiation | | Total time | Total light dose |
|---|---|---|---|---|
| | 405 nm<br>3.6 kW/cm² | 488 nm<br>2.6 kW/cm² | | |
| **Padron** | 60.8 ms | 207 ms | 267.8 ms | 0.76 kJ/cm² |
| **Padron2** | 5.4 ms | 73.1 ms | 78.5 ms | 0.21 kJ/cm² |
| **Kohinoor** | 164.5 ms | 124.9 ms | 289.4 ms | 0.92 kJ/cm² |

**Supplemental Table 2.** Primers used for amplification of the Padron2 coding sequence during cloning of mammalian expression plasmids.

| Target structure/compartment | Primer orientation | Sequence |
|---|---|---|
| vimentin, mitochondria | fw | TCCACCGGTCGCCACCATGGTGAGCAAGGGCGAGGAG |
| vimentin, keratin, mitochrondria | rev | CCCTGCGGCCGCTTTACTTGTACAGCTCGTCCATGGC |
| keratin | fw | GACGGTACCGCGGGCCCGGGATCCACCGGTCGCCACCATGGTGAGCAAGGGCGAGGAG |
| lifeact | fw | AGGGGATCCACCGGTCGCCACCGTGAGCAAGGGCGAGGAGAACAAC |
| lifeact | rev | CGAGCGGCCGCTACTTGTACAGCTCGTCCATGG |
| Map2, CenpC | fw | GATCCGCTAGCGCTAATGGTGAGCAAGGGCGAGGAG |
| Map2, peroxisomes | rev | CACTCGAGATCTGAGTCCGGACTTGTACAGCTCGTCCATGGC |
| ER | fw | CTGCAGGTCGACATGGTGAGCAAGGGCGAGGA |
| ER | rev | TTCTGCGGCCGCCTTGTACAGCTCGTCCATGGCCTGCCCC |
| cytosolic, caveolin | fw | TCCACCGGTCGCCACCATGGTGAGCAAGGGCGAG |
| cytosolic, caveolin | rev | GTCGCGGCCGCTTACTTGTACAGCTCGTC |
| Nup50, histone H2bn | fw | TCCGCTAGCGCTACCGGTCGCCACCATGGTGAGCAAGGGCG |
| Nup50, histone H2bn | rev | CCACTCGAGATCTGAGTCCGGACTTGTACAGCTCGTCCATG |
| CenpC | rev | CACTCGAGATCTGAGTCCGGACTTGTACAGCTCGTCCATG |
| peroxisomes | fw | CGACGCTAGCATGGTGAGCAAGGGCG |
